# Chromosome-level genome assembly of the greenfin horse-faced filefish (*Thamnaconus septentrionalis*) using Oxford Nanopore PromethION sequencing and Hi-C technology

**DOI:** 10.1101/798744

**Authors:** Li Bian, Fenghui Li, Jianlong Ge, Pengfei Wang, Qing Chang, Shengnong Zhang, Jie Li, Changlin Liu, Kun Liu, Xintian Liu, Xuming Li, Hongju Chen, Siqing Chen, Changwei Shao, Zhishu Lin

## Abstract

The greenfin horse-faced filefish, *Thamnaconus septentrionalis*, is a valuable commercial fish species that is widely distributed in the Indo-West Pacific Ocean. It has characteristic blue-green fins, rough skin and spine-like first dorsal fin. *T. septentrionalis* is of a conservation concern as a result of sharply population decline, and it is an important marine aquaculture fish species in China. The genomic resources of this filefish are lacking and no reference genome has been released. In this study, the first chromosome-level genome of *T. septentrionalis* was constructed using Nanopore sequencing and Hi-C technology. A total of 50.95 Gb polished Nanopore sequence were generated and were assembled to 474.31 Mb genome, accounting for 96.45% of the estimated genome size of this filefish. The assembled genome contained only 242 contigs, and the achieved contig N50 was 22.46 Mb, reaching a surprising high level among all the sequenced fish species. Hi-C scaffolding of the genome resulted in 20 pseudo-chromosomes containing 99.44% of the total assembled sequences. The genome contained 67.35 Mb repeat sequences, accounting for 14.2% of the assembly. A total of 22,067 protein-coding genes were predicted, of which 94.82% were successfully annotated with putative functions. Furthermore, a phylogenetic tree was constructed using 1,872 single-copy gene families and 67 unique gene families were identified in the filefish genome. This high quality assembled genome will be a valuable genomic resource for understanding the biological characteristics and for facilitating breeding of *T. septentrionalis*.

## 1 Introduction

The greenfin horse-faced filefish (*Thamnaconus septentrionalis*; hereafter “filefish”) belongs to the family Monacanthidae (Tetraodontiformes) and has characteristic blue-green fins, rough skin and spine-like first dorsal fin (Figure 1)(Su & Li, 2002). It is widely distributed in the Indo-West Pacific Ocean, ranging from the Korean Peninsula, Japan and China Sea to East Africa. Filefish is a temperate demersal species inhabiting a depth range of 50-120 m, and feeding on planktons such as copepods, ostracods, and amphipods, as well as mollusks and benthic organisms(Su & Li, 2002). It goes through annual long-distance seasonal migrations and has diurnal vertical migration habits during wintering and spawning(Lin, Gan, Zheng, & Guan, 1984; Su & Li, 2002). Due to a high protein content and good taste, filefish is an important commercial species in China, Korea and Japan. An interesting feature of filefish is its rough skin, whose roughness is actually attributed to the covered dense small scales. These scales are difficult to remove, and people have to peel off the skin before eating. Given this, filefish is also called “skinned fish” in China.

**FIGURE 1.**
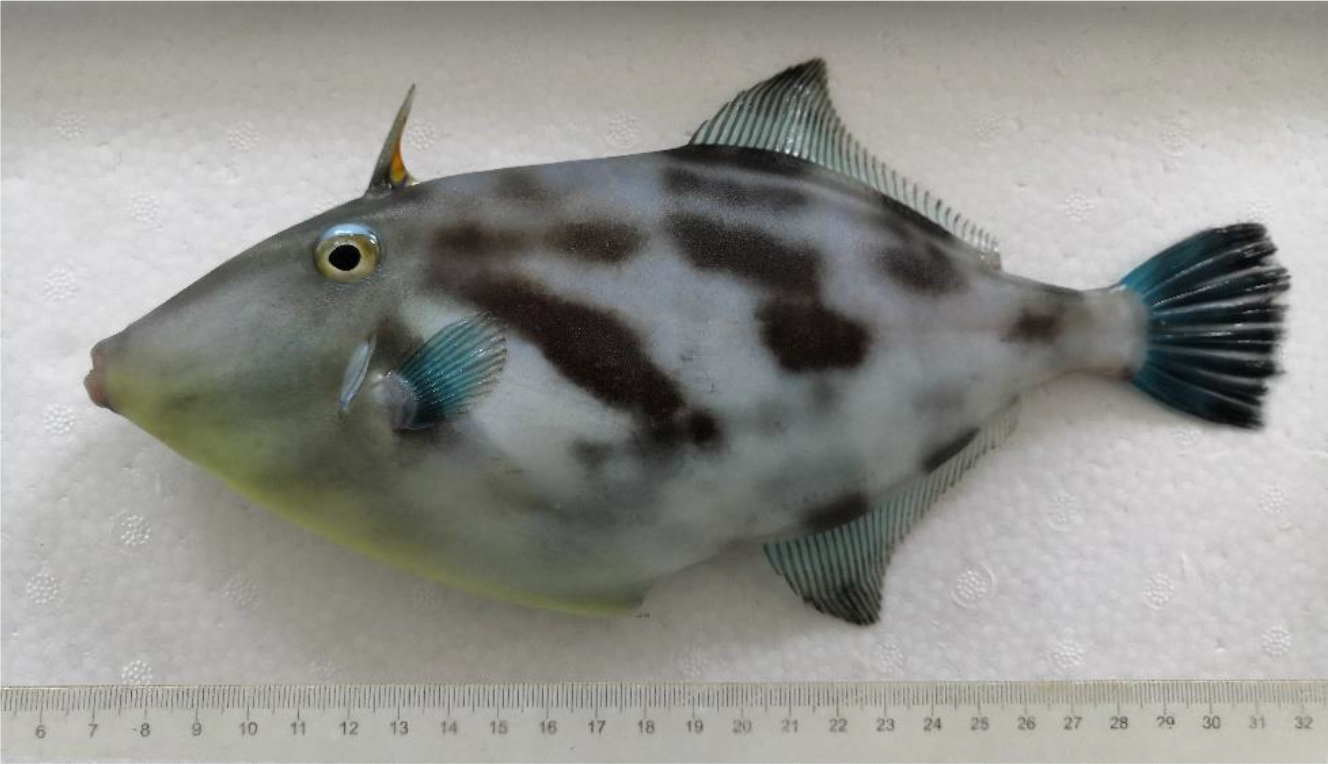
The greenfin horse-faced filefish (*Thamnaconus septentrionalis*)

The wild resource of filefish has declined dramatically since 1990 due to overfishing, and the annual catch in the East China Sea was only 3,842 tons in 1994(Chen, Li, & Hu, 2000). Since then, researchers have attempted to explore the methods to properly culture filefish. Several key technologies including fertilized eggs collection, sperm cryopreservation, larval rearing, tank and cage culturing have been studied, and this species is cultivated commercially in China, Korea and Japan(Guan et al., 2013; Kang et al., 2004; Li, Jiang, Xu, & Liu, 2002; Liu et al., 2017; Mizuno, Shimizu-Yamaguchi, Miura, & Miura, 2012). The current main challenge of filefish cultivation is the high mortality of fish fry during artificial breeding. A better understanding of the underlying genomic-level characteristics will provide significant information to break through the bottleneck and benefit the cultivation industry of this filefish. However, the available genetic information of filefish is scarce. At present, only limited genetic studies regarding microsatellite loci isolation and population structure are available for this filefish (An et al., 2011; An, Lee, Park, & Jung, 2013; Bian et al., 2018; Xu, Chen, & Tian, 2010; Xu, Tian, Liao, & Chen, 2009).

Spectacular improvements in high-throughput sequencing technology, especially the single-molecule sequencing methods, have remarkably reduced the sequencing costs, making a genome project affordable for individual labs. Oxford Nanopore sequencing technology is currently the most powerful method for rapid generation of long-read sequences and has the potential to offer relatively low-cost genome sequencing of non-model animals. It directly detects the input DNA without PCR amplification or synthesis, so the length of sequenced DNA can be very long. The longest read generated by Nanopore sequencing has been up to 2,272,580 bases(Payne, Holmes, Rakyan, & Loose, 2018). Nanopore sequencing has been used in several fish species to construct high-quality genome assembly or to improve the completeness of previous genome drafts(Austin et al., 2017; Ge et al., 2019; Jansen et al., 2017; Kadobianskyi, Schulze, Schuelke, & Judkewitz, 2019; Tan et al., 2018). In the case of red spotted grouper (*Epinephelus akaara*), a chromosome-level reference genome with a contig N50 length of 5.25 Mb was constructed by taking advantage of Nanopore sequencing and Hi-C technology(Ge et al., 2019). In clown anemonefsh (*Amphiprion ocellaris*), a hybrid Illumina/Nanopore method generated much longer scaffolds than Illumina-only approach with an 18-fold increase in N50 length and increased the genome completeness by an additional 16%(Tan et al., 2018).

In this study, the first chromosome-level genome of filefish was constructed using Nanopore sequencing and Hi-C technology. This genomic data will benefit a comprehensive conservation study of filefish along the China and Korea coast to implement better protection of wild populations, and allow us to screen for genetic variations correlated with fast-growth and disease-resistance traits of filefish in the future.

## 2 Materials and methods

### 2.1 Sample and DNA extraction

A single female fish (~325 g) was collected on August 2018 from the Tianyuan Fisheries Co., Ltd (Yantai, China).The muscle tissue below the dorsal fin was taken and stored in the liquid nitrogen until DNA extraction. Genomic DNA was extracted using CTAB (Cetyltrimethylammonium bromide) method. The quality and concentration of the extracted genomic DNA was checked using 1% agarose gel electrophoresis and a Qubit fluorimeter (Invitrogen, Carlsbad, CA, USA). This high-quality DNA was used for subsequent Nanopore and Illumina sequencing.

### 2.2 Library construction and genome sequencing

To generate Oxford Nanopore long reads, approximately 15 μg of genomic DNA was size-selected (30–80 kb) with a BluePippin (Sage Science, Beverly, MA, USA), and processed according to the Ligation Sequencing Kit 1D (SQK-LSK109) protocol. Briefly, DNA fragments were repaired using the NEBNext FFPE Repair Mix (New England Biolabs). After end-reparation and 3’-adenylation with the NEBNext End repair/dA-tailing Module reagents (New England Biolabs), the Oxford Nanopore sequencing adapters were ligated using NEBNext Quick Ligation Module (E6056) (New England Biolabs). The final library was sequenced on 3 different R9.4 flow cells using the PromethION DNA sequencer (Oxford Nanopore, Oxford, UK) for 48 hours. The MinKNOW software (version 2.0) was used to conduct base calling of raw signal data and convert the fast5 files into fastq files. These raw data was then filtered to remove short reads (<5 kb) and the reads with low-quality bases and adapter sequences.

Illumina sequencing libraries were prepared to carry out genome size estimation, correction of genome assembly, and assembly evaluation. The paired-end (PE) libraries with insert sizes of 300 bp were constructed according to the Illumina standard protocol (San Diego, CA, USA) and subjected to PE (2 × 150 bp) sequencing on an Illumina HiSeq X Ten platform (Illumina, San Diego, CA, USA). After discarding the reads with low-quality bases, adapter sequences, and duplicated sequences, the clean reads were used for subsequent analysis.

### 2.3 Genome size estimation and genome assembly

A k-mer depth frequency distribution analysis of the Illumina data was conducted to estimate the genome size, heterozygosity, and content of repetitive sequences of the filefish. The k-mer analysis was carried out using “kmer freq stat” software (developed by Biomarker Technologies Corporation, Beijing, China). Genome size (G) was estimated based on the following formula: G = k-mer number/average k-mer depth, where k-mer number = total k-mers—abnormal k-mers (with too low or too high frequency).

For genome assembly, Canu (version 1.5) (Koren et al., 2017)was conducted for initial read correction, and the assembly was performed by Wtdbg (https://github.com/ruanjue/wtdbg). The consensus assembly was generated by 2 rounds of Racon (version 1.32)(Vaser, Sović, Nagaranjan, & Šikić, 2017), and 3 rounds of Pilon (version 1.21)(Walker et al., 2014) polishing using the Illumina reads with default settings.

### 2.4 Hi-C library construction and sequencing

For Hi-C sequencing, the muscle tissue of filefish was used for library preparation according to Rao et al(Rao et al., 2014). Briefly, the tissue cells were fixed with formaldehyde and restriction endonuclease Hind III was used to digest DNA. The 5’ overhang of the fragments were repaired and labeled using biotinylated nucleotides, followed by ligation in a small volume. After reversal of crosslinks, ligated DNA was purified and sheared to a length of 300-700 bp. The DNA fragments with interaction relationship were captured with streptavidin beads and prepared for Illumina sequencing. The final Hi-C libraries were sequenced on an Illumina HiSeq X Ten platform (Illumina, San Diego, CA, USA) to obtain 2 × 150 bp paired-end reads. To assess the quality of Hi-C data, the plot of insert fragments length frequency was first made to detect the quality of Illumina sequencing. Second, we used BWA-MEM (version 0.7.10-r789) (Li & Durbin, 2009)to align the PE clean reads to the draft genome assembly. In the end, HiC-Pro (Servant et al., 2015) (version 2.10.0) was performed to find the valid reads from unique mapped read pairs.

### 2.5 Chromosomal-level genome assembly using Hi-C data

We first performed a preassembly for error correction of contigs by breaking the contigs into segments of 500 kb on average and mapping the Hi-C data to these segments using BWA-MEM (version 0.7.10-r789)(Li & Durbin, 2009). The corrected contigs and valid reads of Hi-C were used to perform chromosomal-level genome assembly using LACHESIS(Burton et al., 2013) with the following parameters: CLUSTER_MIN_RE_SITES=22; CLUSTER_MAX_LINK_DENSITY=2; CLUSTER_NONINFORMATIVE_RATIO=2; ORDER_MIN_N_RES_IN_TRUNK=10; ORDER_MIN_N_RES_IN_SHREDS=10. To evaluate the quality of the chromosomal-level genome assembly, a genome-wide Hi-C heatmap was generated by ggplot2 in R package.

### 2.6 Assessment of the genome assemblies

To assess the genome assembly completeness and accuracy, we first aligned the Illumina reads to the filefish assembly using BWA-MEM (version 0.7.10-r789)(Li & Durbin, 2009). Furthermore, CEGMA (version 2.5) (Parra, Bradnam, & Korf, 2007)was conducted to find core eukaryotic genes (CEGs) in the genome with parameter set as identity>70%. Finally, the completeness of the genome assembly was also evaluated by using BUSCO (version 2.0)(Simao, Waterhouse, Ioannidis, Kriventseva, & Zdobnov, 2015) search the genome against the actinopterygii database, which consisted of 4584 orthologs.

### 2.7 Repeat annotation, gene prediction and gene annotation

We first used MITE-Hunter(Han & Wessler, 2010), LTR-FINDER (version 1.05)(Xu & Wang, 2007), RepeatScout (version 1.0.5)(Price, Jones, & Pevzner, 2005) and PILER(Edgar & Myers, 2005) to construct a *de novo* repeat library for filefish with default settings. These predicted repeats were classified using PASTEClassifer (version 1.0)(Hoede et al., 2014), and then integrated with Repbase (19.06)(Bao, Kojima, & Kohany, 2015) to build a new repeat library for final repeat annotation. In the end, RepeatMasker (version 4.0.6)(Tarailo-Graovac & Chen, 2009) was performed to detect repetitive sequences in the filefish genome with the following parameters: “-nolow -no_is -norna -engine wublast”.

*Ab initio*-based, homolog-based, and RNA-sequencing (RNA-seq)-based methods were conducted in combination to detect the protein-coding genes in filefish genome assembly. Genscan(Burge & Karlin, 1997), Augustus (version 2.4)(Stanke & Waack, 2003), GlimmerHMM (version 3.0.4)(Majoros, Pertea, & Salzberg, 2004), GeneID (version 1.4)(Blanco, Parra, & Guigó, 2007), and SNAP (version 2006-07-28)(Korf, 2004) were used for *ab initio*-based gene prediction in filefish genome assembly. For the homolog-based method, tiger pufferfish (*Takifugu rubripes*), spotted green pufferfish (*Tetraodon nigroviridis*) and zebrafish (*Danio rerio*) were chosen to conduct gene annotation using GeMoMa (version 1.3.1)(Keilwagen et al., 2016). For the RNA-seq-based method, a mixture of 10 tissues (including brain, eye, gill, heart, liver, intestine, spleen, ovary, kidney and muscle) of a female and the testis of a male filefish was used to construct Illumina sequencing library and subjected to PE (2 × 150 bp) sequencing on an Illumina HiSeq X Ten platform (Illumina, San Diego, CA, USA). After discarding the reads with low-quality bases, adapter sequences, and duplicated sequences, the retained high-quality clean reads were first assembled by Hisat (version 2.0.4)(Kim, Langmead, & Salzberg, 2015) and Stringtie (version 1.2.3)(Pertea et al., 2015), and then the gene prediction was performed using TransDecoder (http://transdecoder.github.io) (version 2.0), GeneMarkS-T (version 5.1)(Tang, Lomsadze, & Borodovsky, 2015), and PASA (version 2.0.2)(Haas et al., 2003). EVM (version 1.1.1)(Haas et al., 2008) was performed to integrate the prediction results obtained from three methods. We then added the genes that were supported by homolog and RNA-seq analysis after-manual evaluation.

To functionally annotate the predicted genes, they were aligned to the Non-redundant protein sequences (NR), eukaryotic orthologous groups of proteins (KOG)(Tatusov et al., 2003), Kyoto Encyclopedia of Genes and Genomes (KEGG) (Kanehisa & Goto, 2000)and TrEMBL(Boeckmann et al., 2003) databases using BLAST (version 2.2.31)(Altschul, Gish, Miller, Myers, & Lipman, 1990) with an e-value cutoff of 1E-5. Gene ontology (GO) (Consortium, 2004)annotation was performed with Blast2GO (version 4.1)(Conesa et al., 2005). For non-coding RNA prediction, we first used tRNAscan-SE (version 1.3.1)(Lowe & Eddy, 1997) to annotate transfer RNAs (tRNAs). Furthermore, Infenal (version 1.1)(Nawrocki & Eddy, 2013) was conducted to search for ribosomal RNAs (rRNAs) and microRNAs based on Rfam (version 13.0)(Daub, Eberhardt, Tate, & Burge, 2015) and miRbase (version 21.0)(Griffiths-Jones, Grocock, Van Dongen, Bateman, & Enright, 2006) database.

### 2.8 Comparative genomics

To resolve the phylogenetic position of the filefish, we first used OrthoMCL (version 2.0.9) (Li, Stoeckert, & Roos, 2003) to detect orthologue groups by retrieving the protein data of eleven teleost species including tiger pufferfish (*Takifugu rubripes*), yellowbelly pufferfish (*Takifugu flavidus*), spotted green pufferfish (*Tetraodon nigroviridis*), red seabream (*Pagrus major*), medaka (*Oryzias latipes*), large yellow croaker (*Larimichthys crocea*), three-spined stickleback (*Gasterosteus aculeatus*), nile tilapia (*Oreochromis niloticus*), japanese seabass (*Lateolabrax maculatus*), spotted gar (*Lepisosteus oculatus*) and zebrafish (*Danio rerio*). The single copy orthologous genes shared by all 12 species were further aligned using MUSCLE (version 3.8.31)(Edgar, 2004) and concatenated to construct a phylogenetic tree with PhyML(Guindon et al., 2010). The divergence time among species was estimated by the MCMCTree program of the PAML package(Yang, 2007) and CAFÉ(version 4.0) (De Bie, Cristianini, Demuth, & Hahn, 2006) was used to identified expanded and contracted gene families.

## 3 Results and discussion

### 3.1 Initial characterization of the filefish genome

The k-mer (k = 19 in this case) depth frequency distribution analysis of the 45.97 Gb clean Illumina data was conducted to estimate the genome size, heterozygosity, and repeat content of filefish (Table 1). The k-mer depth of 76 was found to be the highest peak in the plot, and a k-mer number of 37,677,330,713 was used to calculate the genome size of filefish (Figure S1). The sequences around k-mer depth of 38 were heterozygous sequences, and k-mer depth more than 153 represented repetitive sequences. The filefish genome size was estimated to be 491.74 Mb, the heterozygosity was approximately 0.35%, and the content of repetitive sequences and guanine-cytosine were about 16.62% and 46.05%, respectively.

**TABLE 1.** Statistics of the sequencing data.

| Types | Method | Sequencing platform | Library size (bp) | Clean data (Gb) | Coverage (×)<sup>†</sup> |
| --- | --- | --- | --- | --- | --- |
| Genome | Illumina | Illumina HiSeq X | 300 | 45.97 | 93.48 |
| Genome | Nanopore | PromethION | ultra-long | 50.95 | 103.61 |
| Genome | Hi-C | Illumina HiSeq X | 300 | 39.44 | 80.20 |
| Transcriptome | Illumina | Illumina HiSeq X | 300 | 11.31 | 23.00 |

<sup>†</sup> The coverage was calculated using an estimated genome size of 491.74 Mb.

### 3.2 Genome assembly

A total of 50.95 Gb high quality clean reads, representing a 104‐fold coverage of the genome, were generated from PromethION DNA sequencer (Table 1, Table S1-2),. These data was assembled using Wtdbg, followed by Racon and Pilon polishing, which produced a 465.93 Mb genome assembly with a surprising long contig N50 of 22.07 Mb (Table S3). The length of this assembly was close to the genome size estimated by k-mer analysis (491.74 Mb), indicating an appropriate assembly size was obtained from the Nanopore data. Among the sequenced tetraodontiform species, the genome size of filefish was larger than *Takifugu* and *Tetraodon* species, but smaller than *Mola mola*(Aparicio et al., 2002; Gao et al., 2014; Jaillon et al., 2004; Pan et al., 2016) (Table 2).

**TABLE 2.** Assembly statistics of filefish and other tetraodontiform genomes.

| Species | *T. septentrionalis* | *Takifugu rubripes*<sup>†</sup> | *Takifugu flavidus* | *Tetraodon nigroviridis* | *Mola mola* |
| --- | --- | --- | --- | --- | --- |
| Sequencing technology | Oxford Nanopore sequencing | PacBio Sequel | PacBio Sequel | Plasmid library + BAC library sequencing | Illumina Hiseq 2000 |
| Assembly size (Mb) | 474.31 | 384.13 | 366.29 | 342.40 | 639.45 |
| Number of scaffolds | 155 | 128 | 867 | 25773 | 5552 |
| N50 scaffold size (Mb) | 23.05 | 16.71 | 15.68 | 0.73 | 8.77 |
| Number of contigs | 242 | 530 | 1111 | 41566 | 51826 |
| N50 contig length (Mb) | 22.46 | 3.14 | 4.36 | 0.03 | 0.02 |

<sup>†</sup> The assembly statistics of other tetraodontiform genomes were from NCBI assembly database. The GenBank assembly accession numbers were as follows: *Takifugu rubripes* (GCA\_901000725.2), *Takifugu flavidus* (GCA\_003711565.2), *Tetraodon nigroviridis* (GCA\_000180735.1), *Mola mola* (GCA\_001698575.1).

For Hi-C data, overall 39.44 Gb clean reads were obtained and used for subsequent analysis (Table 1). To assess the quality of Hi-C data, we first made a plot of insert fragments length frequency, which showed a relatively narrow unimodal length distribution with the highest peak around 350 bp (Figure S2), indicating efficient purification of streptavidin beads during library construction. The alignment results revealed that about 89.78% of the Hi-C read pairs were mapped on the genome, and 78.18% of the read pairs were unique detected on the assembly (Table S4). Lastly, a total of 47,111,219 valid reads, which accounted for 66.95% of the unique mapped reads, were detected by HiC-Pro in the Hi-C dataset (Table S5). Taken together, our evaluation suggested an overall high quality of the Hi-C data, and only the valid read pairs were used for subsequent analysis.

Before chromosomal-level genome assembly, an error correction of the initial assembly was performed by BWA-MEM with Hi-C data. The corrected filefish genome assembly was approximately 474.30 Mb with only 242 contigs, the contig N50 reached up to 22.46 Mb, and the longest contig was 32.32 Mb (Table 2, Table S6). The results indicated that high-coverage Nanopore long read-only assembly, followed by multiple iterations of genome polishing using Illumina reads is an effective method to generate high-quality genome assemblies.

A chromosomal-level genome was then assembled using LACHESIS, the results showed that overall 147 contigs spanning 471.65 Mb (99.44% of the assembly) were scaffolded into 20 pseudo-chromosomes, and 107 contigs spanning 469.46 Mb (98.98% of the assembly) were successfully ordered and oriented (Table 3). Several of the pseudo-chromosomes were scaffolded with only 2 or 3 contigs, representing a high contiguity of the genome. The final assembled genome was 474.31 Mb with a scaffold N50 length of 23.05 Mb and a longest scaffold of 34.81 Mb (Table 2, Table S6). As far as we know, this assembled genome was one of the most contiguous fish genome assembly with the highest contig N50 when compared with other published fish genomes.

**TABLE 3.** Statistics of the pseudo-chromosome assemblies using Hi-C data.

| Group | Contig number | Contig length (bp) |
| --- | --- | --- |
| Group 1 | 3 | 34,805,468 |
| Group 2 | 3 | 34,142,503 |
| Group 3 | 3 | 29,239,029 |
| Group 4 | 13 | 27,092,115 |
| Group 5 | 3 | 24,789,104 |
| Group 6 | 7 | 24,144,372 |
| Group 7 | 10 | 23,815,151 |
| Group 8 | 3 | 23,107,901 |
| Group 9 | 11 | 22,985,309 |
| Group 10 | 5 | 23,048,615 |
| Group 11 | 2 | 22,982,431 |
| Group 12 | 6 | 23,025,906 |
| Group 13 | 3 | 22,547,364 |
| Group 14 | 11 | 22,005,842 |
| Group 15 | 16 | 20,921,416 |
| Group 16 | 3 | 20,603,809 |
| Group 17 | 2 | 19,738,352 |
| Group 18 | 5 | 17,694,734 |
| Group 19 | 13 | 18,094,054 |
| Group 20 | 25 | 16,862,837 |
| Total contigs clustered | 147 | 471,646,312 |
| Total contigs ordered and oriented | 107 | 469,464,378 |

To further evaluate the quality of the chromosomal-level genome assembly, a genome-wide Hi-C heatmap was generated. The 20 pseudo-chromosomes could be easily distinguished and the interaction signal strength around the diagonal was much stronger than that of other positions within each pseudo-chromosome, which indicated a high quality of this genome assembly (Figure 2).

**FIGURE 2.**
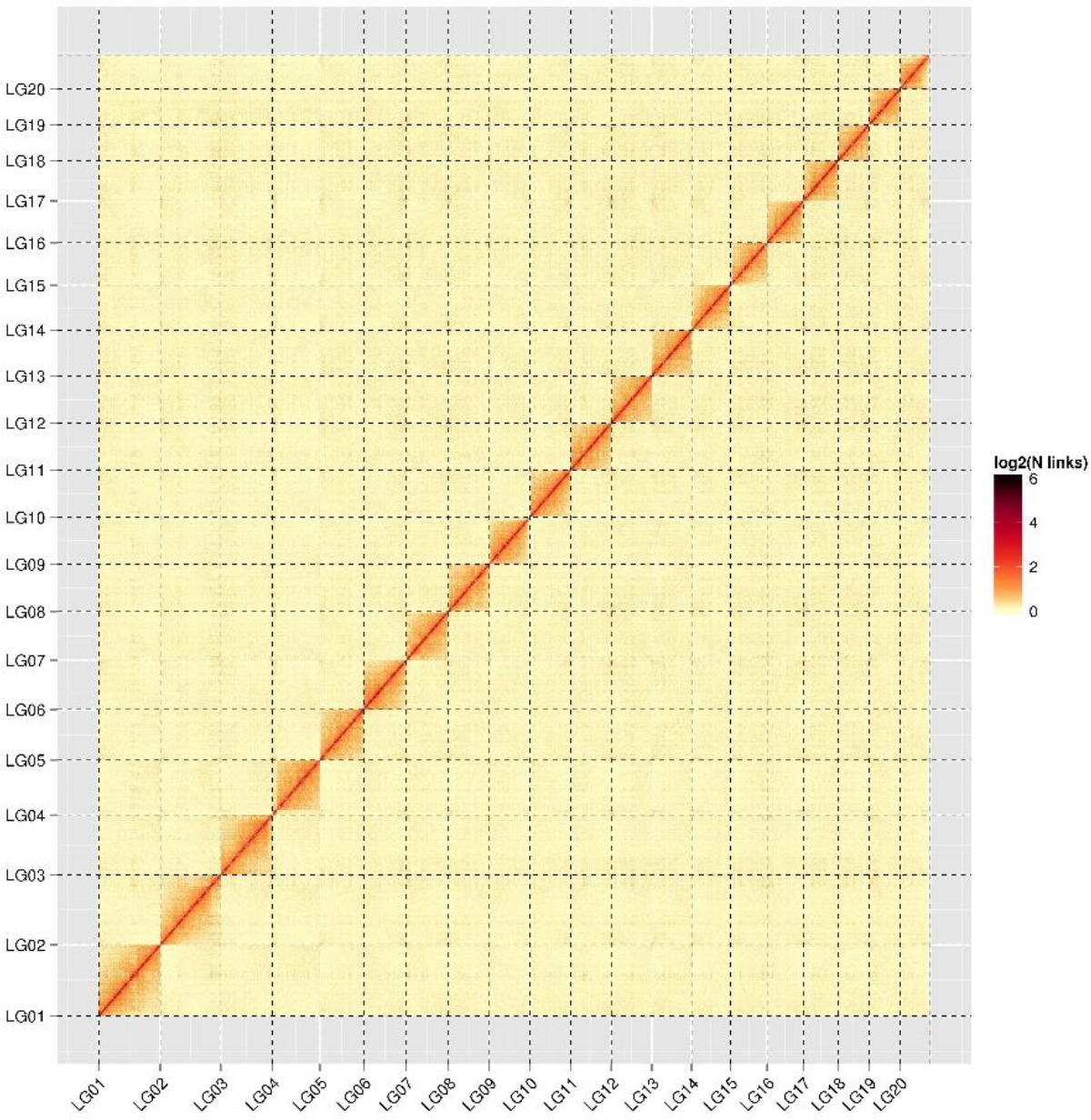
The genome-wide Hi-C heatmap of the filefish. LG 1-20 are the abbreviations of Lachesis Group 1-20, representing the 20 pseudo-chromosomes.

### 3.3 Completeness of the assembled genome

Illumina reads were aligned to the filefish assembly, and 97.41% of the clean reads can be mapped to the contigs (Table S7). Then the CEGMA analysis identified 442 CEGs, accounting for 96.51% of all 458 CEGs in the program, and 226 CEGs could be detected by using a highly conserved 248 CEGs dataset (Table S8). Lastly, approximately 94.33% (4324/4584) of complete BUSCOs were found in the assembly (Table S9). Overall, the assessment results indicated our filefish genome assembly was complete and of high quality.

### 3.4 Repeat annotation, gene prediction and gene annotation

A total of 67.35 Mb of repeat sequences that accounted for 14.2% of the assembly were found in filefish (Table S10). This repeat content was close to the value (16.62%) obtained from k-mer analysis. The predominant repeats type were TIRs (4.35%), LINEs (2.40%) and LARDs (1.65%).

The combination of *Ab initio*-based, homolog-based, and RNA-seq-based methods predicted overall 22,067 protein-coding genes with an average gene length, average exon length, and average intron length of 11,291bp, 230 bp, and 905 bp, respectively (Table 1, Table 4). A total of 20,924 genes, which counted for 94.82% of the predicted genes, were successfully annotated with putative functions (Table 5). The non-coding RNA prediction identified 1,703 tRNAs, 649 rRNAs and 109 microRNAs, respectively (Table S11).

**TABLE 4.**
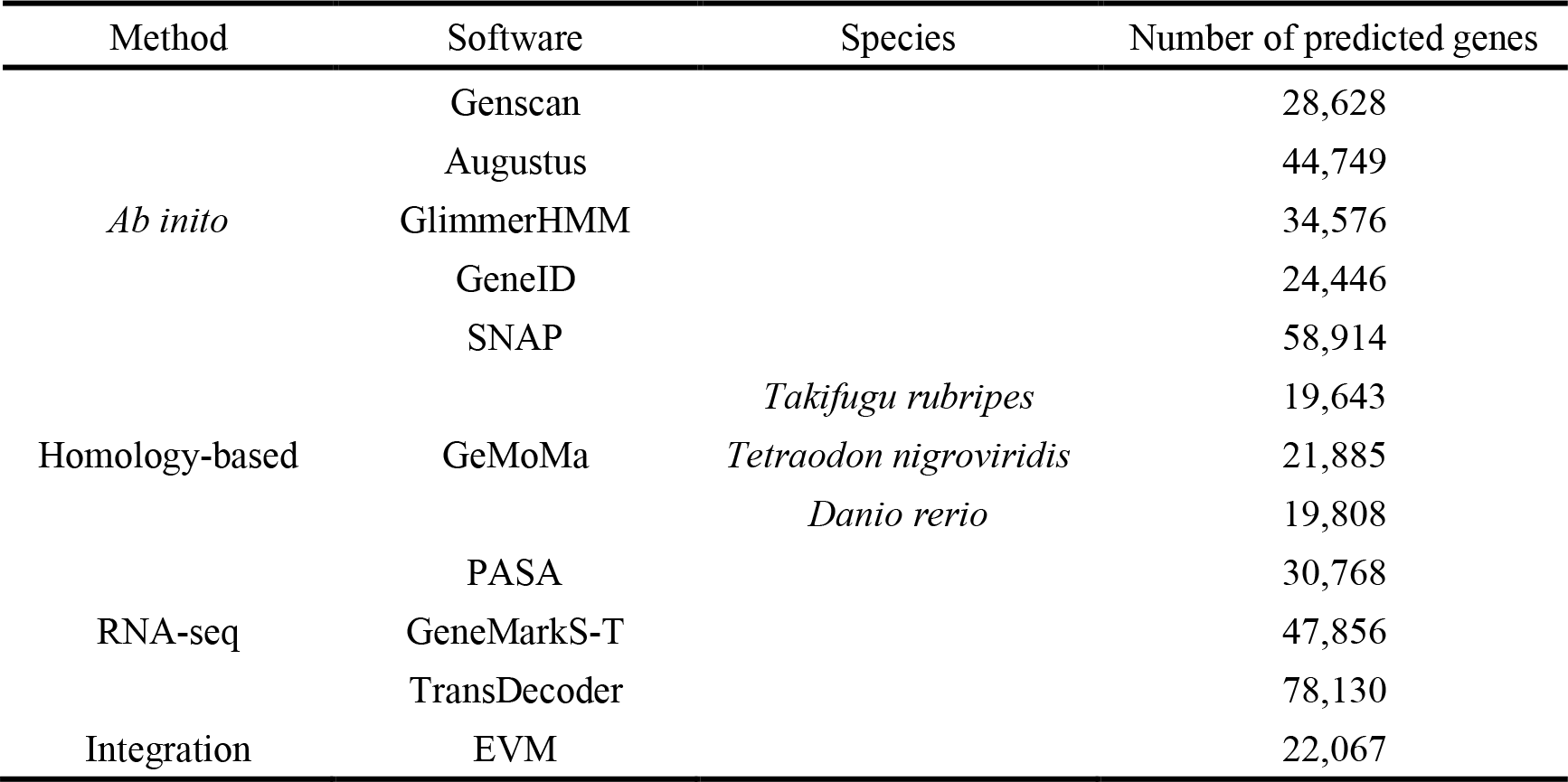
Summary of predicted protein-coding genes in the filefish genome.

**TABLE 5.** Summary of functional annotations for predicted genes.

| Annotation database | Annotated number of predicted genes | Percentage (%) |
| --- | --- | --- |
| GO | 11,257 | 51.01% |
| KEGG | 13,714 | 62.15% |
| KOG | 14,760 | 66.89% |
| TrEMBL | 20,795 | 94.24% |
| NR | 20,905 | 94.73% |
| All Annotated | 20,924 | 94.82% |
| Predicted Genes | 22,067 | - |

### 3.5 Comparative genomics

Comparison of the filefish genome assembly with other eleven teleost species genomes found a total of 22,665 gene families, of which 5,692 were shared among all eleven species, including 1,872 single‐copy orthologous genes (Table S12). Overall 20,261 genes of filefish can be clustered into 15,433 gene families, including 67 unique gene families containing 193 genes (Table S12). The phylogenetic tree showed that four tetraodontiform species were clustered together, and the divergence time between filefish and the other three species was around 124.4 million years ago (Mya) (Figure 3). We also found 59 expanded gene families and 98 contracted gene families in filefish compared with the other fish species (Figure S3). A Venn diagram of orthologous gene families among four tetraodontiform species was also constructed, and 971 unique gene families containing 6485 genes were identified in the filefish genome (Figure 4).

**FIGURE 3.**
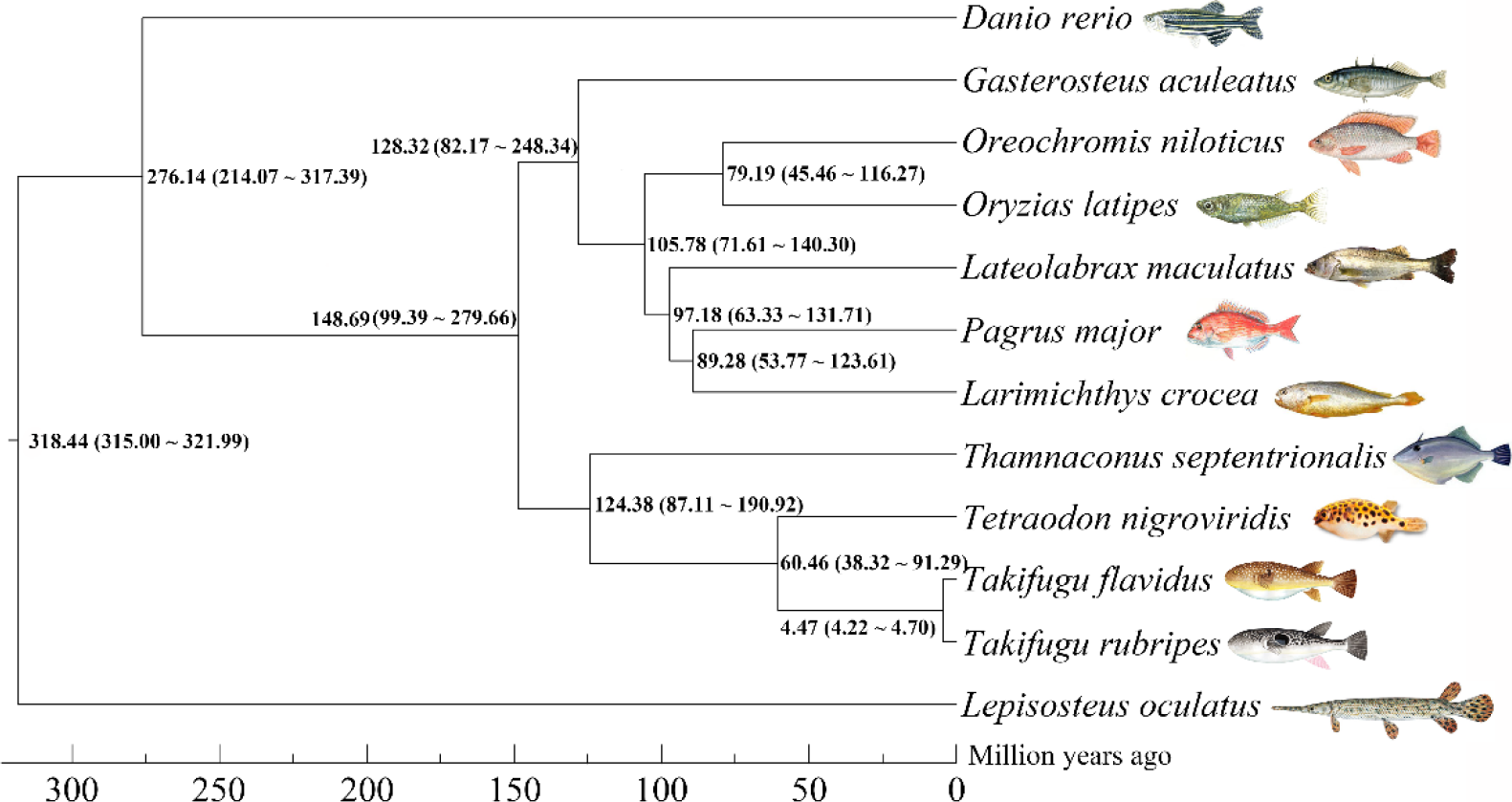
Phylogenetic analysis of the filefish with other teleost species. *Lepisosteus oculatus* was used as the outgroup. The estimated species divergence time (million years ago) and the 95% confidential intervals were labeled at each branch site.

**FIGURE 4.**
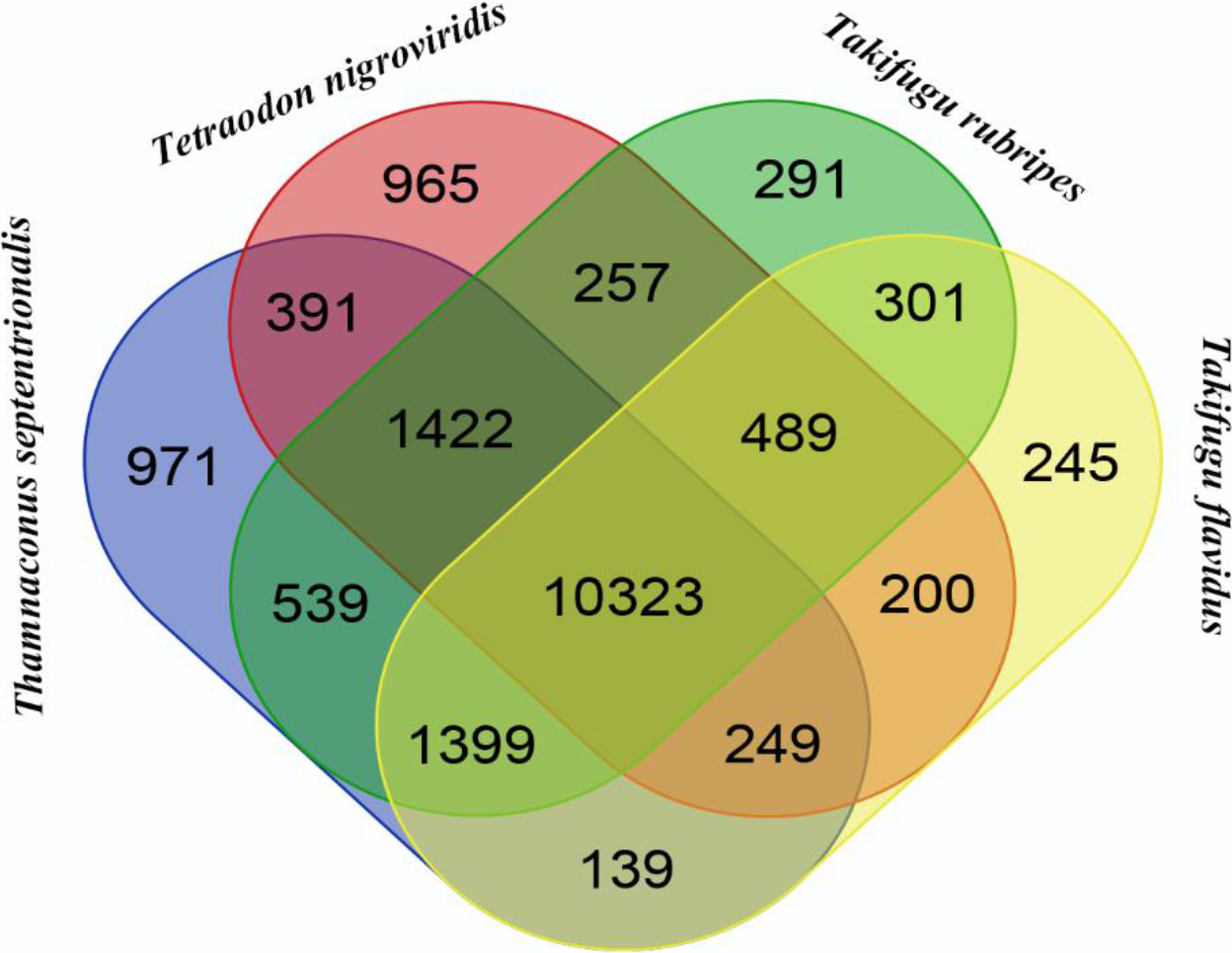
Venn diagram of orthologous gene families among four tetraodontiform species.

## 4 Conclusion

In the present study, we assembled the chromosome-level genome of *T. septentrionalis*, a first reference genome of the genus *Thamnaconus*. The assembled genome was 474.31 Mb, which is larger than the sequenced *Takifugu* and *Tetraodon* species, but smaller than *Mola mola*. With the powerful sequencing ability of Oxford Nanopore technology, the contig N50 of the assembled genome achieved 22.46 Mb, and the longest contig was 32.32 Mb. To the best of our knowledge, this is the highest contig N50 among all the sequenced fish genomes. This revealed that a combination of high-coverage Nanopore sequencing and Illumina data polishing can effectively produce highly contiguous genome assemblies. The contigs were clustered and ordered onto 20 pseudo-chromosomes with Hi-C data, and several pseudo-chromosomes were scaffolded with only 2 or 3 contigs. This high-quality genome will lay a strong foundation for a range of breeding, conservation and phylogenetic studies of filefish in the future.

## Acknowledgements

We appreciate the help from Tianyuan Fisheries Co., Ltd (Yantai, China) who provided the filefish samples. This work was supported by fund of Key Laboratory of Open-Sea Fishery Development, Ministry of Agriculture, P. R. China (LOF 2017-05), fund of Guangdong Provincial Key Laboratory of Fishery Ecology and Environment, South China Sea Fisheries Research Institute, Chinese Academy of Fisheries Sciences, SCSFRI, CAFS (FEEL-2017-10), Key Research and Development Program of Shandong province, Department of Science & Technology of Shandong province (2019GHY112073) and Central Public-interest Scientific Institution Basal Research Fund, YSFRI, CAFS (20603022017014).

## Data Accessibility

Raw sequencing reads are available on GenBank as BioProject PRJNA565600. Raw sequencing data (Nanopore, Illumina, Hi-C and RNA-seq data) have been deposited in SRA (Sequence Read Archive) database as SRX6875837, SRX6862879, SRX6875660, and SRX6875519.

## Author Contributions

S.C., C.S. and Z.L. designed and managed the project. L.B., F.L. and J.G. interpreted the data and drafted the manuscript. P.W., S.Z., C.L. and X.L. prepared the materials. Q.C., J.L., K.L. and H.C. preformed the DNA extraction, RNA extraction and libraries construction. L.B., F.L., X.L. and C.S. performed the bioinformatic analysis. All authors contributed to the final manuscript editing.

