## Supplemental Tables and Figures for "Chromosome-level genome assembly of the greenfin horse-faced filefish (*Thamnaconus septentrionalis*) using Oxford Nanopore PromethION sequencing and Hi-C technology"

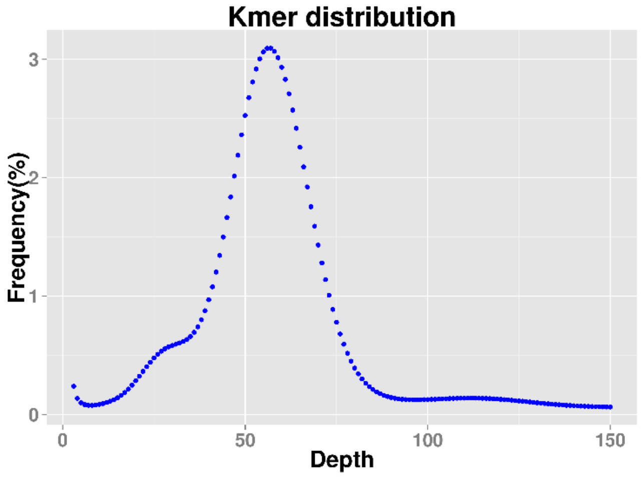


**FIGURE S1 Distribution of k-mers of length 19 from the Illumina data**

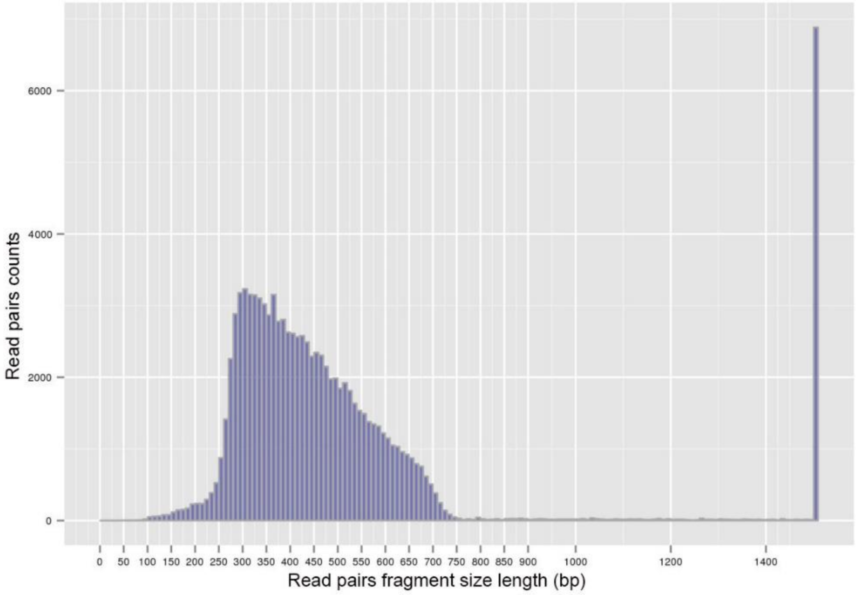


**FIGURE S2 The length distribution of the insert fragments**

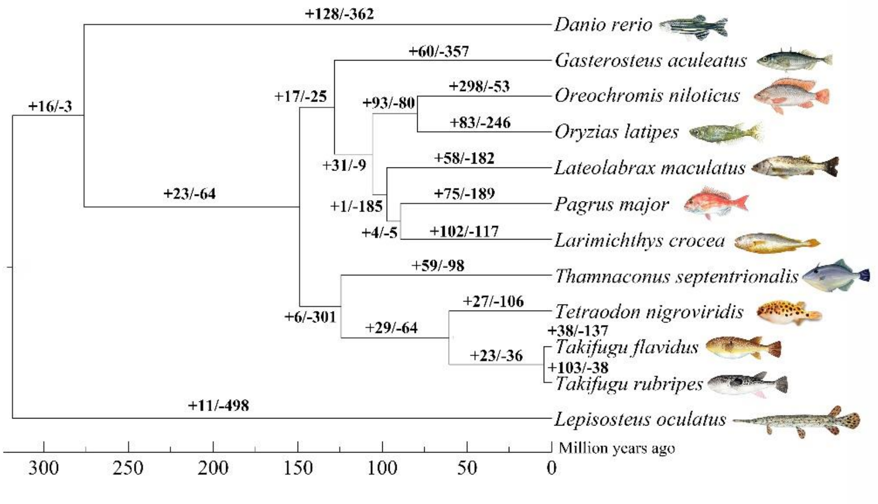


**FIGURE S3 Phylogenetic relationship of the filefish and other teleost species. The number of expanded gene families (positive values) and number of contracted gene families (negative values) were labeled at each branch.**

**TABLE S1 The Nanopore subreads used for genome assembly**

| Number | Total length (bp) | N50 length (bp) | Average length (bp) |
|---|---|---|---|
| 3,325,617 | 50,951,959,068 | 20,946 | 15,321 |

**TABLE S2 Size distribution of the Nanopore subreads**

| Length (bp) | Number | Total length (bp) | Average length (bp) |
|---|---|---|---|
| 5000~10000 | 1,681,760 | 11,742,329,829 | 6,982 |
| 10000~20000 | 926,472 | 12,866,908,175 | 13,888 |
| 20000~30000 | 327,016 | 7,998,097,285 | 24,458 |
| 30000~40000 | 177,722 | 6,126,543,930 | 34,473 |
| 40000~50000 | 93,774 | 4,172,932,320 | 44,500 |
| 50000~60000 | 51,713 | 2,819,834,600 | 54,529 |
| 60000~70000 | 28,678 | 1,851,327,652 | 64,556 |
| 70000~80000 | 16,513 | 1,230,644,062 | 74,526 |
| 80000~ | 21,969 | 2,143,341,215 | 97,562 |
| Total | 3,325,617 | 50,951,959,068 | 15,321 |

**TABLE S3 Statistics of the filefish assembly before Hi-C correction**

| Contig number | Contig length (bp) | Contig N50 (bp) | Contig N90 (bp) | Contig max (bp) | GC content (%) | Gap total length (bp) |
|---|---|---|---|---|---|---|
| 233 | 465,933,797 | 22,068,396 | 14,714,403 | 31,748,062 | 45.54 | 0 |

**TABLE S4 The alignment of the Hi-C clean data to the genome assembly**

| Mapping Type | Number | Ratio (%) |
|---|---|---|
| Total Read Pairs | 90,000,000 | 100 |
| Mapped Reads | 161,607,721 | 89.78 |
| Unique Mapped Read Pairs | 70,364,370 | 78.18 |

**TABLE S5 Statistics of different types of read pairs produced by Hi-C sequencing**

| Type | Number | Ratio (%) |
|---|---|---|
| Unique paired alignments | 70,364,370 | 100 |
| Valid interaction pairs | 47,111,219 | 66.95 |
| Dangling end pairs | 11,804,913 | 16.78 |
| Re-ligation pairs | 4,956,284 | 7.04 |
| Self-cycle pairs | 484,581 | 0.69 |
| Dumped pairs | 6,007,373 | 8.54 |

**TABLE S6 Statistics of the Hi-C assembly of the filefish genome**

| Statistical level | Scaffold | Contig |
|---|---|---|
| Total number | 155 | 242 |
| Total length (bp) | 474,309,635 | 474,300,935 |
| N50 Length (bp) | 23,049,015 | 22,457,894 |
| N90 Length (bp) | 17,560,257 | 14,961,343 |
| Maximum length (bp) | 34,805,668 | 32,321,032 |

**TABLE S7 The alignment of the Illumina reads to the filefish genome assembly**

| Total reads | Mapped reads | Mapped (%) |
|---|---|---|
| 306,820,704 | 298,862,082 | 97.41 |

**TABLE S8 CEGMA assessment of the filefish genome assembly**

| Number of 458 CEGs present in assembly | Percentage of 458 CEGs present in assembly (%) | Number of 248 highly conserved CEGs present in assembly | Percentage of 248 highly conserved CEGs present in assembly (%) |
|---|---|---|---|
| 442 | 96.51 | 226 | 91.13 |

**TABLE S9 BUSCO assessment of the filefish genome assembly**

| Complete BUSCOs | Complete single-copy BUSCOs | Complete duplicated BUSCOs | Fragmented BUSCOs | Missing BUSCOs |
|---|---|---|---|---|
| 4,324 (94.33%) | 4,213 (91.91%) | 111 (2.42%) | 62 (1.35%) | 198 (4.32%) |

**TABLE S10 Statistics of the repeat sequences in the filefish genome**

| Type | Number | Length (bp) | Percentage (%) |
|---|---|---|---|
| ClassI/DIRS | 2,483 | 263,934 | 0.06 |
| ClassI/LINE | 81,085 | 11,388,389 | 2.40 |
| ClassI/LTR | 43,384 | 4,826,403 | 1.02 |
| ClassI/LTR/Copia | 1,282 | 192,552 | 0.04 |
| ClassI/LTR/Gypsy | 28,575 | 3,741,163 | 0.79 |
| ClassI/PLE\|LARD | 52,693 | 7,838,084 | 1.65 |
| ClassI/SINE | 5,584 | 857,903 | 0.18 |
| ClassI/SINE\|TRIM | 180 | 32,195 | 0.01 |

| | | | |
|---|---|---|---|
| ClassI/TRIM | 6,007 | 3,084,453 | 0.65 |
| ClassI/Unknown | 2,955 | 248,514 | 0.05 |
| ClassII/Crypton | 1,622 | 137,731 | 0.03 |
| ClassII/Helitron | 12,313 | 940,637 | 0.20 |
| ClassII/MITE | 25,140 | 5,195,286 | 1.10 |
| ClassII/Maverick | 11,144 | 808,998 | 0.17 |
| ClassII/TIR | 190,397 | 20,616,935 | 4.35 |
| ClassII/Unknown | 83,453 | 6,686,885 | 1.41 |
| PotentialHostGene | 3,758 | 485,772 | 0.10 |
| Total | 552,055 | 67,345,834 | 14.21 |

**TABLE S11 Statistics of the non-coding RNA in the filefish genome**

| RNA classification | Number | Family |
|---|---|---|
| tRNA | 1,703 | 25 |
| rRNA | 649 | 4 |
| miRNA | 109 | 21 |

**TABLE S12 Statistics of the gene families of the twelve teleost species**

| Species | Number of total genes | Number of one copy genes | Number of multi-copy genes | Number of unigene | Number of other genes | Number of clustered genes | Number of unclustered genes | Number of total gene families | Number of unique gene families |
|---|---|---|---|---|---|---|---|---|---|
| *T. septentrionalis* | 22,067 | 3,818 | 5,720 | 193 | 10,530 | 20,261 | 1,806 | 15,433 | 67 |
| *T.flavidus* | 20,707 | 3,526 | 6,849 | 73 | 8,663 | 19,111 | 1,596 | 13,345 | 36 |
| *T.rubripes* | 20,996 | 3,707 | 6,083 | 166 | 10,149 | 20,105 | 891 | 15,021 | 50 |
| *T.nigroviridis* | 27,918 | 3,722 | 6,039 | 672 | 9,138 | 19,571 | 8,347 | 14,296 | 255 |
| *D.rerio* | 32,258 | 2,917 | 9,412 | 4,091 | 14,026 | 30,446 | 1,812 | 15,572 | 741 |
| *P.major* | 28,343 | 3,959 | 4,906 | 1,570 | 9,593 | 20,028 | 8,315 | 14,489 | 508 |
| *O.latipes* | 22,039 | 3,745 | 5,940 | 728 | 11,083 | 21,496 | 543 | 15,470 | 146 |
| *L.crocea* | 23,283 | 3,583 | 6,491 | 325 | 12,166 | 22,565 | 718 | 16,577 | 99 |
| *G.aculeatus* | 20,770 | 3,755 | 5,819 | 476 | 9,471 | 19,521 | 1,249 | 14,187 | 57 |
| *O.niloticus* | 29,504 | 3,573 | 6,789 | 3,167 | 14,743 | 28,272 | 1,232 | 16,849 | 471 |
| *L.maculatus* | 21,622 | 4,070 | 4,648 | 682 | 9,089 | 18,489 | 3,133 | 13,935 | 237 |
| *L.oculatus* | 18,717 | 4,369 | 3,643 | 613 | 8,251 | 16,876 | 1,841 | 13,568 | 156 |
